# NKG2A-mediated immune modulation of natural killer cells by *Staphylococcus aureus*

**DOI:** 10.1101/2024.11.29.625999

**Authors:** Kate Davies, Al-Motaz Rizek, Simon Kollnberger, Eddie C. Y. Wang, Matthias Eberl, Jonathan Underwood, James E. McLaren

## Abstract

Natural killer (NK) cells are specialized lymphocytes that help protect against viruses and cancer. However, in the context of bacterial infections, NK cells can be harmful, rather than protective. Such immune pathogenesis by NK cells has been linked to the over-production of pro-inflammatory cytokines like interferon-γ (IFN-γ). In this context, IFN-γ-deficient mice display increased survival rates in response to *Staphylococcus aureus* (*S. aureus*), which causes life-threatening, invasive systemic infections with high mortality rates in humans. However, little is known about how NK cells respond to *S. aureus* in humans. In this study, we found that the peripheral blood of patients with bloodstream *S. aureus* infection was enriched for NKG2A^+^ NK cells with greater cytokine producing capacity, compared to those hospitalized with *Escherichia coli* bloodstream infections. As a possible mechanistic cause, superantigens from *S. aureus* promoted the expansion of CD57^−^ NKG2A^+^ NK cells which produced IFN-γ through an IL-12-independent mechanism and exhibited reduced levels of CD16 compared to unstimulated NK cells. These data suggest that *S. aureus* bloodstream infection in humans promotes a phenotypic shift towards NKG2A^+^ NK cells with greater IFN-γ producing capacity, providing a plausible way to promote inflammation-driven disease pathogenesis.

**AUTHOR SUMMARY:** Natural Killer (NK) cells are specialized immune cells that provide crucial defence against viruses but can also respond to bacterial infections, especially in humans. During bloodstream infection by *Staphylococcus aureus*, a Gram-positive bacterial pathogen that causes life-threatening infections in humans, NK cells may actually be harmful, rather than protective, to the host. However, very little is known about the NK cell response to invasive *Staphylococcus aureus* infections in humans. Here, we show that human patients with bloodstream *Staphylococcus aureus*, but not *Escherichia coli*, infections have an increased frequency of NK cells with increased pro-inflammatory capacity. Furthermore, we show that toxins produced by *Staphylococcus aureus*, which help the bacteria evade the protective effects of T cells the immune system (“superantigens”), promoted the expansion of these pro-inflammatory NK cells, providing a possible mechanistic cause for their increased presence in patients with bloodstream *Staphylococcus aureus* infections. Collectively, these results suggest that *Staphylococcus aureus* infection triggers phenotypic and functional changes in NK cells that provide a plausible way to promote inflammation-driven disease pathogenesis

## INTRODUCTION

*Staphylococcus aureus* is an opportunistic, Gram-positive bacterium which commonly exists as a commensal organism in humans asymptomatically colonising specific anatomical sites, such as the nasal mucosa(1), skin and gastrointestinal tract(2). Such residency requires the bacterium to overcome inbuilt host defence mechanisms, notably those orchestrated by innate (e.g. neutrophils) and adaptive (e.g. T cells) immune cells which provide major forms of protective immunity(3). This commensalism is a risk factor for the human host to develop life-threatening, invasive infections such as bloodstream infections (or “bacteraemia”) and endocarditis(4), which are associated with high mortality rates (30-day mortality at 20-30%)(5, 6). *S. aureus* is one of the most frequent causes of community- or hospital-acquired bacteraemia, second only to the Gram-negative bacterium *Escherichia coli* (*E. coli*)(7). *S. aureus* is also a leading cause of sepsis(8), a life-threatening condition caused by an exaggerated host immune response to infection that is estimated to underlie 1 in 5 deaths worldwide(9). Morbidity and mortality are exacerbated by its ability to establish deep-seated, metastatic infections despite seemingly effective antimicrobial therapy (10).

The inherent threat from *S. aureus* bacteraemia highlights the need to understand the pathophysiological interactions between this bacterium and the human immune system, including the immune evasive strategies that promote its persistence and survival. In this regard, *S. aureus* utilises a complex arsenal of virulence factors and immune evasion proteins that prevent immunological elimination by neutrophils, macrophages, T cells and antibodies(11). These include pore-forming toxins, such as leukocidin ED (LukED), Panton-Valentine leukocidin and α-hemolysin, which bind to and directly kill neutrophils, macrophages and/or T cells, and highly potent “superantigen” (SAg) exotoxins, which act to counteract and weaken host T cell defence mechanisms(12–14). *S. aureus* is capable of producing 26 distinct SAgs encoded with mobile genetic elements including toxic shock syndrome (TSS) toxin-1 (TSST-1) and staphylococcal enterotoxins (SEs), such as SEA, SEB, and SEC1-3(15, 16). Distinct *S. aureus* strains can encode multiple SAgs, although rarely produce >20 different SAgs at the same time (15, 17). SAgs primarily function by cross-linking T cell receptors on αβ T cells to major histocompatibility complex (MHC) class II molecules on the surface of antigen-presenting cells(11, 14, 16). Such engagement causes SAgs to overstimulate T cells, leading to excessive cytokine release that may contribute to the propagation of a cytokine “storm” whilst simultaneously driving the upregulation of co-inhibitory receptors on T cells that renders them hyporesponsive(14, 18–21). Consequently, SAgs can drive life-threatening complications of *S. aureus* infection, such as TSS, necrotizing pneumonia and infective endocarditis(15, 22). SAgs can also bind costimulatory molecules(23–26), cytokine receptors(27) and extracellular matrix proteins(28) on the surface of different cell types, providing additional mechanisms of action for immune evasion. SAgs have therefore become targets of therapeutic intervention(29, 30) in an era of increasing antimicrobial resistance (e.g. methicillin-resistant *S. aureus*(31)) and the absence of a licenced vaccine against *S. aureus*.

Natural killer (NK) cells are innate immune cells of lymphocyte origin that play a specialist role in protecting the host against infectious pathogens and malignancy(32). Unlike T cells and B cells, NK cells lack somatically rearranged antigen receptors to mediate their function. Instead, NK cells utilize an array of germline-encoded activating and inhibitory receptors to recognize cell surface molecules of self or foreign origin, such as the killer cell immunoglobulin-like receptors (KIRs), natural cytotoxicity receptors (e.g. NKp30, NKp46), and the C-type lectin NKG2 receptors (e.g. NKG2A, NKG2C, NKG2D)(33). Human NK cells typically emerge into the circulation from secondary lymphoid organs as CD3^−^ CD56^hi^ CD16^−^ cells(34) expressing high levels of NKp30, NKG2A and receptors for pro-inflammatory cytokines (e.g. interleukin-12 (IL-12), IL-18)(35). These CD56^hi^ NK cells possess superior cytokine-producing capacity, despite having limited cytotoxic potential, but can differentiate into CD3^−^ CD56^dim^ CD16^+^ subsets that predominate in the circulation and undergo phenotypic changes to specialize their function towards receptor-driven cytotoxicity(32, 36). These include expression of KIRs, CD57 and the activating NKG2C receptor whilst simultaneously losing NKG2A/CD94 expression(36–38). These highly differentiated NK cells can home to infected or dysregulated tissue sites to mediate their function, whilst many human organs retain populations of tissue-resident NK cells expressing established markers of residency(39–41). NK cell responses to acute or resolving viral infections are typically cytokine-driven, with less differentiated NK cells predominantly responding to IL-12, IL-15, IL-18 and type I interferons (IFNs) produced during infection which promote their survival, proliferation and effector function (e.g. through IFN-γ production)(35, 42, 43). However, in response to chronic infections, such as human cytomegalovirus (HCMV), more differentiated NK cells expand and develop adaptive features, akin to T cells, such as clonal expansion and the formation of immunological memory(44, 45). These adaptive-like NK cell expansions typically express NKG2C and CD57(46, 47). NK cells are essential for antiviral protection since patients with inborn NK cell deficiencies are highly prone to severe or recurrent herpesvirus infections(48).

In contrast to our knowledge of NK cell-mediated immunity to viruses, little is known about how NK cells respond to bacterial infections, especially in humans. NK cells can be activated by bacteria, directly through Toll-like receptors(49) or indirectly by pro-inflammatory cytokines produced by bacteria-sensing antigen presenting cells (e.g. dendritic cells)(50, 51). NK cell-deficient mice have increased bacterial burden and poor immune control in response to *Legionella pneumophilia*(52) and *Mycobacterium tuberculosis*(53). Furthermore, NK cell-produced IFN-γ helps to promote bacterial elimination during various infections, including *Klebsiella pneumoniae*(52–54), whilst memory-like NK cells mediate protective immune responses to *Streptococcus pneumoniae*(55) and *Ehrlichia muris*(56). In contrast, NK cell-deficient or depleted mice have improved survival rates after *E. coli* or *Streptococcus pyogenes* infection, which has been linked to reduced levels of inflammation and lower production of pro-inflammatory cytokines, including IL-12 and IFN-γ(57, 58). Furthermore, NK cells are thought to have a detrimental role in sepsis, including in immunocompromised hosts(59, 60). This points to important pathogen-specific differences in the control and immunopathology of bacterial infections by NK cells. In the context of *S. aureus* infection, NK cells are in fact the major source of IFN-γ (61, 62), and IFN-γ^−/−^ mice or those treated with depleting anti IFN-γ antibodies are more resistant to lethal *S. aureus* challenge(63, 64). The fact that SAgs(65, 66) and other virulence factors such as the pore-forming toxins LukED(67), α-hemolysin(68) and β-hemolysin(69) alter the functionality of NK cells, including increasing their production of IFN-γ, suggests that NK cell-released IFN-γ could be a driver of immune pathogenesis during *S. aureus* infection. However, studies of NK cell responses to systemic *S. aureus* infections have been confined to animal models.

To address this knowledge gap, we here examined the NK cell response to *S. aureus* or *E. coli* bacteraemia in humans. We found that more polyfunctional CD56^dim^ NK cells with greater cytokine producing potential were observed in patients with *S. aureus* bacteraemia, compared to those hospitalized with *E. coli* bacteraemia. These highly responsive NK cells were enriched for NKG2A^+^ subsets. In support of a possible underlying mechanism, SAgs, namely TSST-1, promoted the *in vitro* expansion of CD57^−^ NKG2A^+^ NK cells that had reduced levels of CD16 and were capable of producing IFN-γ through an IL-12-independent mechanism. These data suggest that systemic *S. aureus* infection in humans promotes a phenotypic and functional shift towards IFN-γ producing NKG2A^+^ NK cells, providing a plausible way to promote inflammation-driven disease pathogenesis.

## RESULTS

### Circulating NKG2A^+^ NK cells increase in frequency *in vivo* during *S. aureus* acteraemia

In order to study the NK cell response to *S. aureus*, we used multi-colour flow cytometry panels to identify and phenotype (Figure S1) NK cell subsets (e.g. CD56^dim^ and CD56^hi^ NK cells) in peripheral blood collected from adult patients hospitalized with *S. aureus* bacteraemia (*n* = 7) (Figure 1A). Peripheral blood samples from adult patients hospitalized with *E. coli* bacteraemia (*n* = 8) and healthy controls (*n* = 8) were collected for comparison (Table 1). When assessing global NK cell frequency, patients with *S. aureus*, but not *E. coli,* bacteraemia had slightly lower frequencies of CD56^dim^ NK cells compared to healthy controls, although these differences were not statistically significant (Figures S2A and S2B). Using NKG2A, NKG2C and CD57 as markers of different NK cell differentiation states, we discovered higher percentages of NKG2C^−^ NKG2A^+^ CD56^dim^ NK cells in patients with *S. aureus* bacteraemia, compared to healthy controls or patients with *E. coli* bacteraemia (Figure 1B, 1C & S3A). This increase in NKG2C^−^ NKG2A^+^ cells in *S. aureus* patients was NK cell-specific as it was not recapitulated in CD56^+^ T cells (Figure 1C & S3B), which includes invariant-like subsets such as mucosal associated invariant T (MAIT) cells (70), where the percentage of NKG2C^−^ NKG2A^+^ CD56^+^ T cells was lower than in healthy controls. Furthermore, we found that, amongst all CD56^dim^ NK cells, there was a slight increase in the percentage of CD57^−^ NKG2A^+^ and CD57^+^ NKG2A^+^ NK cells in patients with *S. aureus* bacteraemia, compared to healthy controls (Figure 1B, 1C & S3C). In contrast, there was a significant decrease in CD57^+^ NKG2A^+^ CD56^+^ T cells in patients with *S. aureus* bacteraemia, compared to healthy controls (Figure 1D & S3D). In support of this subtle phenotypic change in NK cells, NKG2A expression levels were higher on CD56^dim^ NK cells from *S. aureus* bacteraemia patients, compared to healthy controls and patients with *E. coli* bacteraemia, but we saw no change in CD57 expression (Figure 1E). Paired analyses of CD56^dim^ NK cells from PBMCs collected at hospital admission and approx. one month after hospital discharge (convalescence) demonstrated that this trend towards more NKG2C^−^ NKG2A^+^ NK cells in *S. aureus* bacteraemia patients was sustained at convalescence, although some patients showed a reduction in the percentage of CD57^−^ NKG2A^+^ NK cells (Figures S3E & S3F). IL- 12 is known to upregulate NKG2A expression on NK cells (71, 72). However, plasma IL-12p70 levels were lower in *S. aureus* bacteraemia patients compared to healthy controls and patients with *E. coli* bacteraemia (Figure S4A), despite a significant positive correlation between plasma IL-12p70 levels and the percentage of NKG2C^−^ NKG2A^+^ NK cells in *S. aureus* patients (Figure S4B).

**Figure 1.**
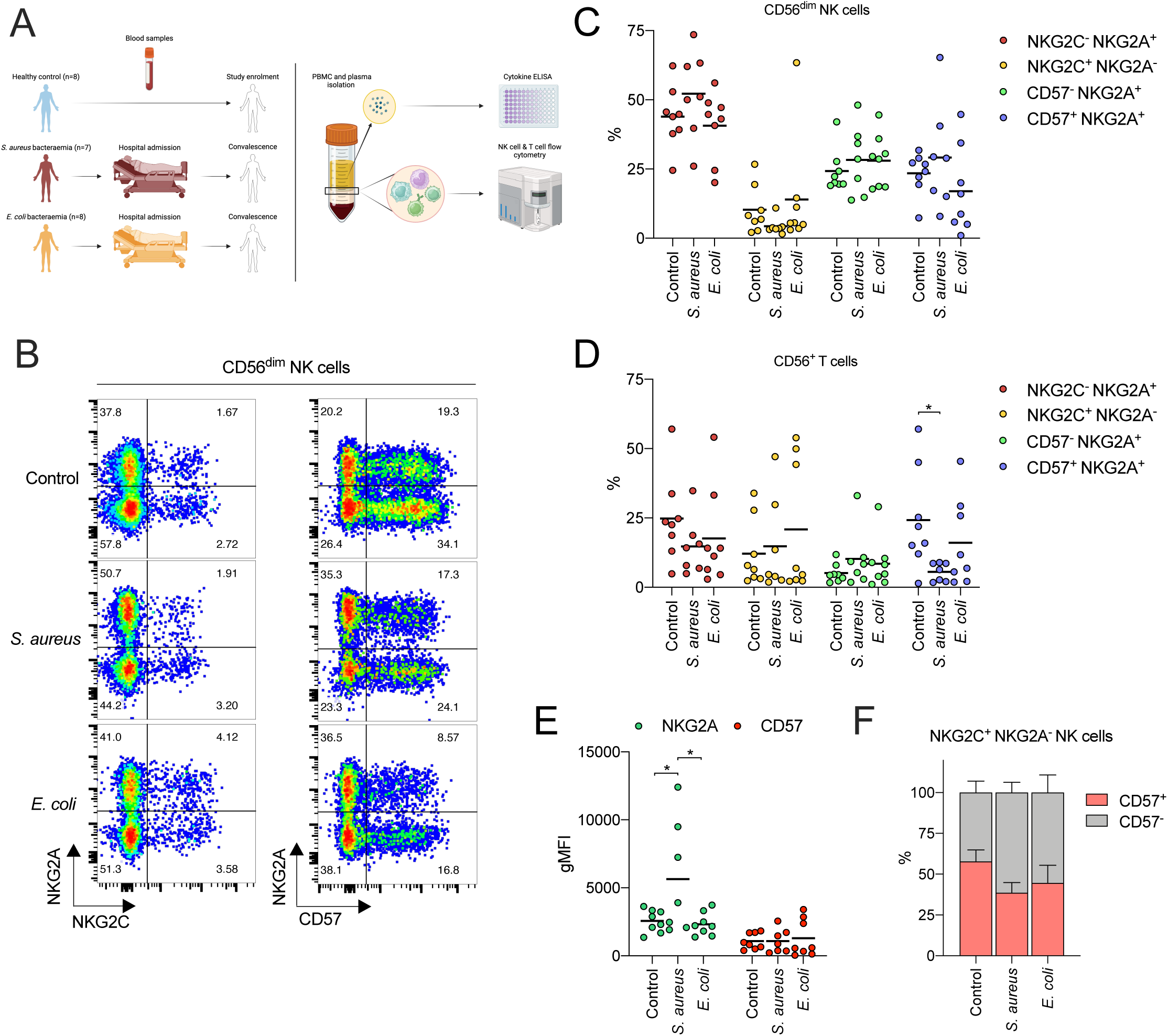
Circulating NKG2A^+^ NK cells expand *in vivo* during *S. aureus* bacteraemia in humans. (A) Schematic overview of study design. (B) Representative flow cytometry plots depicting expression levels of NKG2A vs NKG2C or NKG2A vs CD57 on CD56^dim^ NK cells from human PBMCs collected from healthy controls and patients hospitalized with *S. aureus* or *E. coli* bacteraemia. (C-D) Scatter dot plot showing the frequency of NKG2C^−^ NKG2A^+^ (red filled circles), NKG2C^+^ NKG2A^−^ (yellow filled circles), CD57^−^ NKG2A^+^ (green filled circles), or CD57^+^ NKG2A^+^ (blue filled circles) subsets among CD56^dim^ NK cells (C) or CD56^+^ T cells (D) from human PBMCs collected from healthy controls and patients hospitalized with *S. aureus* or *E. coli* bacteraemia. Each dot represents one control or patient. Line indicates mean value. *p<0.05; One-way ANOVA with Tukey’s post-hoc test. (E) Scatter dot plots depicting geometric mean fluorescence intensity (gMFI) of NKG2A (green filled circles) or CD57 (red filled circles) expression on CD56^dim^ NK cells from human PBMCs collected from healthy controls and patients hospitalized with *S. aureus* or *E. coli* bacteraemia. Each dot represents one donor. Line indicates mean value. **p < 0.05; One-way ANOVA with Tukey’s post-hoc test. (F) Stacked bar charts showing the frequency of CD57^+^ or CD57^−^ cells amongst NKG2C^+^ NKG2A^−^ NK cells from human PBMCs collected from healthy controls (*n =* 8) and patients hospitalized with *S. aureus* (*n =* 7) or *E. coli* (*n =* 8) bacteraemia. Data are shown as mean ± SEM.

**Table 1.**
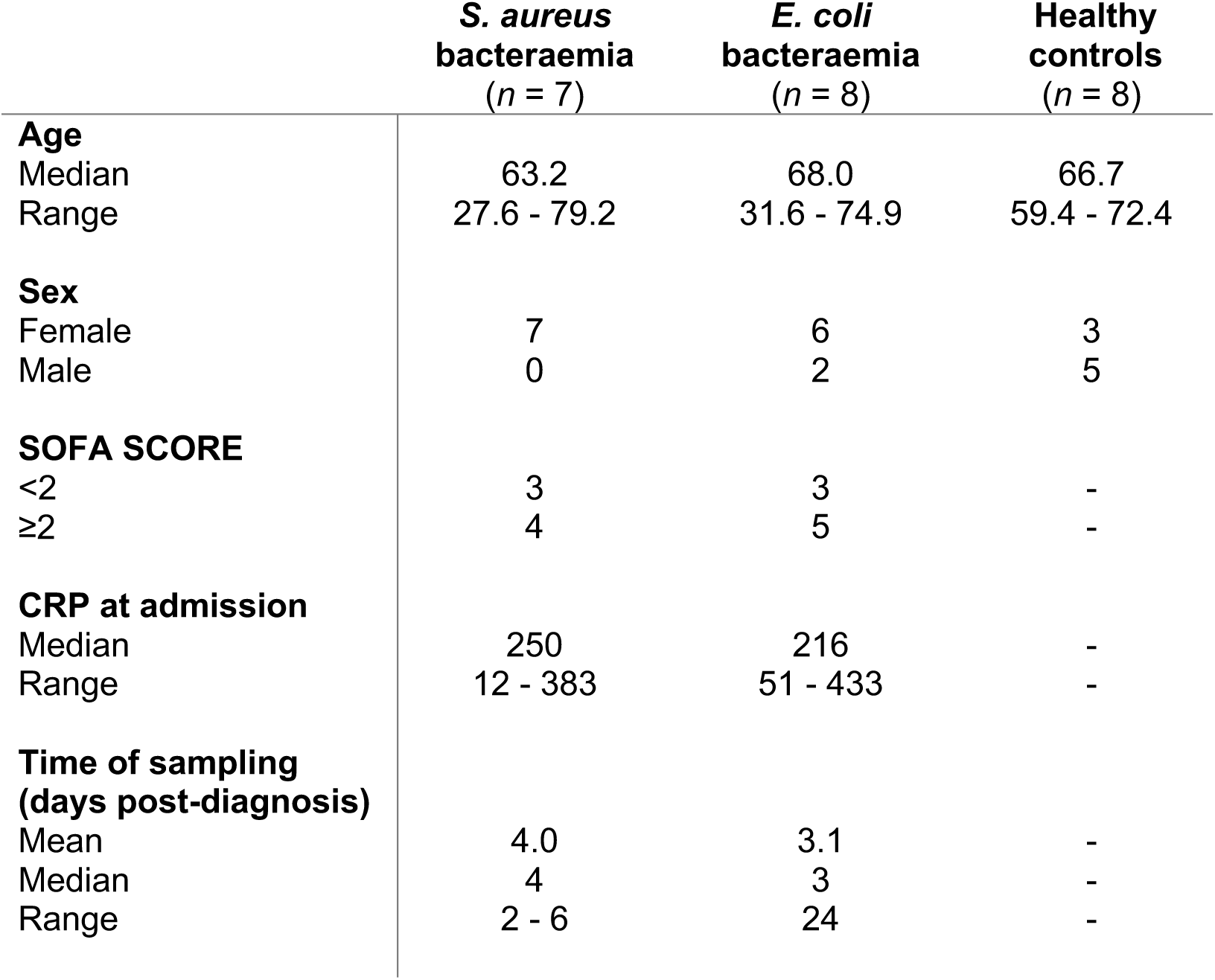
Baseline demographics of adult patients hospitalized with *S. aureus* or *E. coli* bacteraemia compared to healthy controls. Abbreviations: SOFA: sequential organ failure assessment. CRP: c-reactive protein. Diagnosis indicates positive blood culture for *S. aureus* or *E. coli*

|  | ***S. aureus***<br>**bacteraemia**<br>**(n = 7)** | ***E. coli***<br>**bacteraemia**<br>**(n = 8)** | **Healthy**<br>**controls**<br>**(n = 8)** |
| --- | --- | --- | --- |
| **Age** |  |  |  |
| Median | 63.2 | 68.0 | 66.7 |
| Range | 27.6 - 79.2 | 31.6 - 74.9 | 59.4 - 72.4 |
| **Sex** |  |  |  |
| Female | 7 | 6 | 3 |
| Male | 0 | 2 | 5 |
| **SOFA SCORE** |  |  |  |
| <2 | 3 | 3 | - |
| ≥2 | 4 | 5 | - |
| **CRP at admission** |  |  |  |
| Median | 250 | 216 | - |
| Range | 12 - 383 | 51 - 433 | - |
| **Time of sampling**<br>**(days post-diagnosis)** |  |  |  |
| Mean | 4.0 | 3.1 | - |
| Median | 4 | 3 | - |
| Range | 2 - 6 | 24 | - |

To further explore the significance of these NKG2C^−^ NKG2A^+^ NK cell-specific alterations in *S. aureus* patients, we assessed their activation status *ex vivo* using CD69 as a marker of NK cell activation. We found that there was an increase in the percentage of CD69^+^ cells amongst NKG2C^−^ NKG2A^+^ NK cells in patients with *S. aureus* bacteraemia, compared to healthy controls (Figure S5A). However, this increase in CD69^+^ cells was similarly observed in NKG2C^−^ NKG2A^+^ NK cells from patients with *E. coli* bacteraemia, but also within NKG2C^+^ NKG2A^−^ and global CD56^dim^ NK cell populations in both patient groups (Figures S5A, S5B & S5C). This suggested that the presence of either bacterial organism was driving a general increase in NK cell activation that was not restricted to NKG2C^−^ NKG2A^+^ NK cells.

In contrast to data on CD69, *S. aureus*, but much less so *E. coli,* patients displayed a lower percentage of NKG2C^−^ NKG2A^+^ NK cells expressing T-cell immunoglobulin and ITIM domain (TIGIT) compared to healthy controls (Figure S5D), a co-inhibitory receptor found on T cells and NK cells that can suppress NK cell-mediated cytokine production and cytotoxicity(73–76). An inverse correlation between the percentage of CD69^+^ and TIGIT^+^ cells was observed on NKG2C^−^ NKG2A^+^ NK cells from controls or patients with *E. coli* bacteraemia. In contrast, in *S. aureus* patients, we identified a significant positive correlation between CD69 and TIGIT expression on NKG2C^−^ NKG2A^+^ NK cells (Figure S5E). This trend in a reduction of TIGIT^+^ cells was also observed amongst NKG2C^+^ NKG2A^−^ NK cells and global CD56^dim^ NK cell populations in patients with *S. aureus* bacteraemia, compared to healthy controls (Figures S5D & S5F). Furthermore, a similar positive correlation between the percentage of CD69^+^ and TIGIT^+^ cells was observed when analysing the global CD56^dim^ NK cell population (Figure S5G), which suggested that this association between CD69 and TIGIT positivity was more specific to the impact of *S. aureus* on all NK cells.

Adaptive-like NK cells typically represent highly differentiated CD57^+^ NKG2C^+^ populations that preferentially lack expression of NKp30 and the inhibitory receptor Siglec-7(46, 47, 77, 78). Our data demonstrated a reduction in NKG2C^+^ NKG2A^−^ and CD57^+^ NKG2A^−^ CD56^dim^ NK cells in *S. aureus* bacteraemia patients (Figures 1C, S3A & S3C) which might suggest that adaptive-like NK cell phenotypes could also be reduced in *S. aureus* bacteraemia patients. Indeed, we found the percentage of CD57^+^ cells amongst NKG2C^+^ NKG2A^−^ CD56^dim^ NK cells to be lower in *S. aureus* bacteraemia patients (Figure 1F). Therefore, we next sought to determine whether NK cells with an adaptive-like phenotype were altered in *S. aureus* bacteraemia patients. After pre-gating on NKp30^−^ Siglec-7^−^ CD56^dim^ NK cells, there was a reduction in the percentage of NKG2C^+^ NKG2A^−^ subsets in patients with *S. aureus* bacteraemia, compared to healthy controls and *E. coli* bacteraemia patients (Figures 2A & 2B). In agreement with our data on all CD56^dim^ NK cells, we also saw the proportion of NKG2C^−^ NKG2A^+^ and CD57^−^ NKG2A^+^ subsets increase in *S. aureus* bacteraemia patients (Figures 2A, 2B & 2C). Furthermore, the proportion of CD57^+^ NKG2C^+^ adaptive-like NK cells was lower in patients with *S. aureus* bacteraemia, compared to healthy controls or *E. coli* patients (Figure 2D).

**Figure 2.**
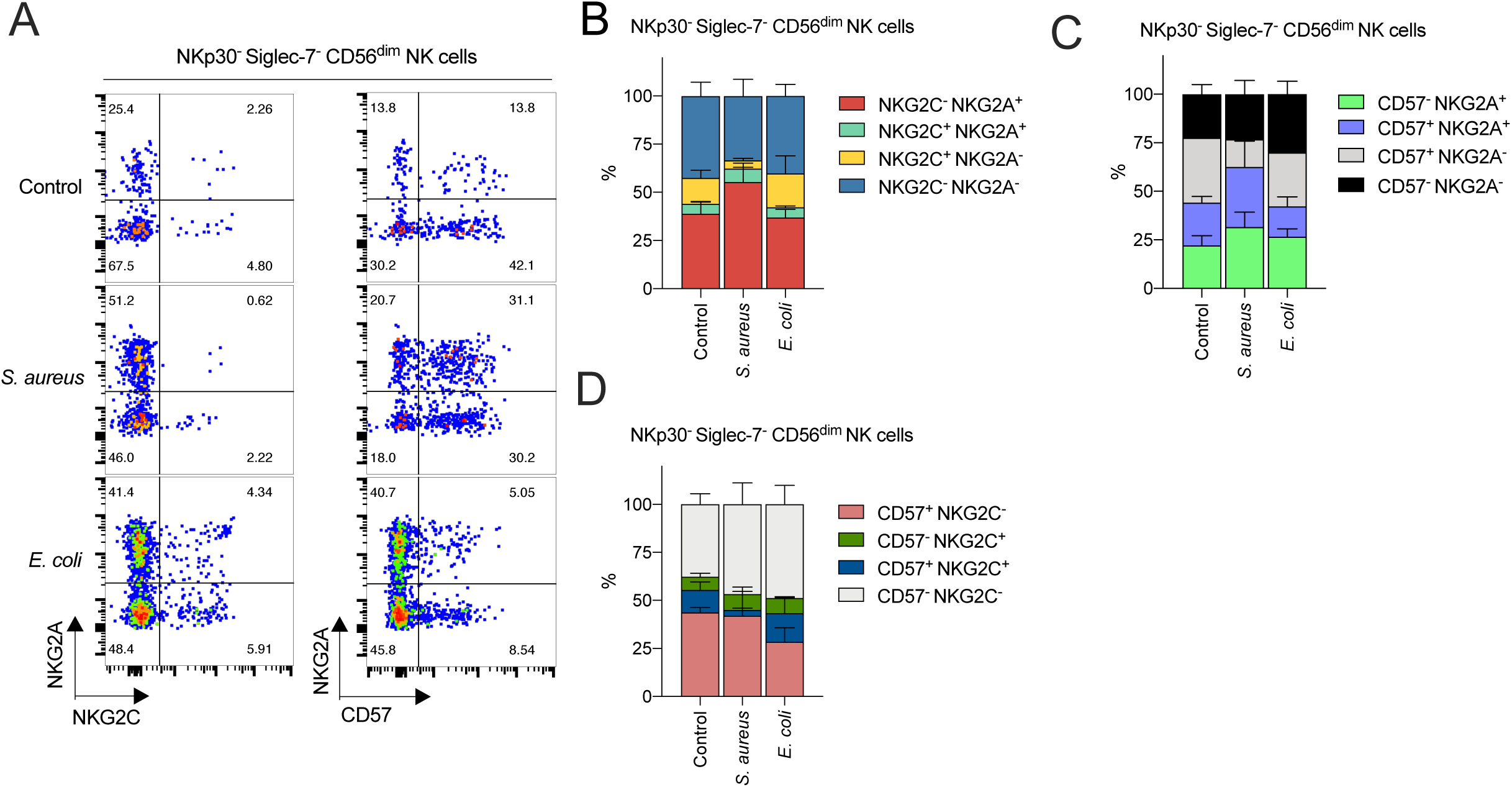
The proportion of CD57^+^ NKG2C^+^ adaptive-like NK cells is lower in patients with *S. aureus* bacteraemia. (A) Representative flow cytometry plots depicting expression levels of NKG2A vs NKG2C or NKG2A vs CD57 on NKp30^+^ Siglec-7^+^ CD56^dim^ NK cells from human PBMCs collected from healthy controls and patients hospitalized with *S. aureus* or *E. coli* bacteraemia. (B-D) Stacked bar charts showing the frequency of NKG2C^−^ NKG2A^+^, NKG2C^+^ NKG2A^+^, NKG2C^+^ NKG2A^−^ and NKG2C^−^ NKG2A^−^ (B), CD57^−^ NKG2A^+^, CD57^+^ NKG2A^+^, CD57^+^ NKG2A^−^, CD57^−^ NKG2A^−^ (C) or CD57^−^ NKG2C^+^, CD57^+^ NKG2C^+^, CD57^+^ NKG2C^−^, CD57^−^ NKG2C^-^ (D) subsets among NKp30^+^Siglec-7^+^ CD56^dim^ NK cells from human PBMCs collected from healthy controls (*n =* 7) and patients hospitalized with *S. aureus* (*n =* 5) or *E. coli* (*n =* 8) bacteraemia. Data are shown as mean ± SEM.

Collectively, these data show that NKG2A^+^ NK cells increase in frequency in patients with *S. aureus* bacteraemia, with a concurrent reduction in the percentage of CD57^+^ NKG2C^+^ adaptive-like NK cells.

### SAgs promote the expansion of CD57^−^ NKG2A^+^ NK cells

Virulence factors encoded by *S. aureus* are capable of targeting NK cells and/or modifying their functionality(65–69). SAgs, which typically target T cells and HLA class II^+^ APCs, can indirectly activate human NK cells, which produce more IFN-γ in response to SAg stimulation than innate-like T cell subsets such as MAIT cells(65, 66). SAgs also promote the production of IL-12 from myeloid cells, which is important for inducing IFN-γ production, rather than cytotoxic granule release, from human NK cells(66, 79, 80). Given that IL-12 also upregulates NKG2A expression on both NK cells(71, 72) and T cells(81), we next examined whether candidate SAgs from *S. aureus* could promote the formation of NKG2A^+^ NK cells *in vitro*. Using PBMCs from healthy donors, stimulation with recombinant forms of SEA, SEB and TSST-1 (24h) produced significant increases in the percentage of IFN-γ^+^, CD107a^+^ and CD69^+^CD56^dim^ NK cells (Figures S6A, S6B, S6C & S6D). Indeed, the degree of activation in CD56^dim^ NK cells was comparable to that seen in CD56^+^ T cells and higher than that seen in more conventional, CD56^−^ T cells, in agreement with published data(66). Importantly though, all three SAgs promoted an increase in the percentage of NKG2C^−^ NKG2A^+^ CD56^dim^ NK cells, but also CD57^−^ NKG2A^+^ populations (Figures 3A, 3B, S7A & S7B). These effects were most evident following TSST-1 stimulation which yielded significant increases in the percentage of NKG2C^−^ NKG2A^+^ and CD57^−^ NKG2A^+^ CD56^dim^ NK cells (Figure 3B) but also in the expression levels of NKG2A on the surface of CD56^dim^ NK cells (Figure 3C). This effect was confined to CD56^dim^ NK cells as all three SAgs yielded very little change in the percentage of NKG2C^−^ NKG2A^+^ CD56^hi^ NK cells, which were already at high levels (∼70%) in unstimulated cells (Figure S7C) whilst the proportion of CD57^−^ NKG2A^+^ cells within CD56^hi^ NK cells even reduced upon stimulation (Figure S7D). The largest proportion of IFN-γ^+^ CD56^dim^ NK cells in response to SAg stimulation was confined to the CD57^−^ NKG2C^−^ NKG2A^+^ subset (Figure 3D & S8). Furthermore, the percentage of TSST-1-induced IFN-γ^+^ cells was significantly higher in CD57^−^NKG2A^+^ CD56^dim^ NK cells, compared to NKG2C^+^ NKG2A^−^ or CD57^+^ NKG2A^+^ NK cell subsets (Figure 3E).

**Figure 3.**
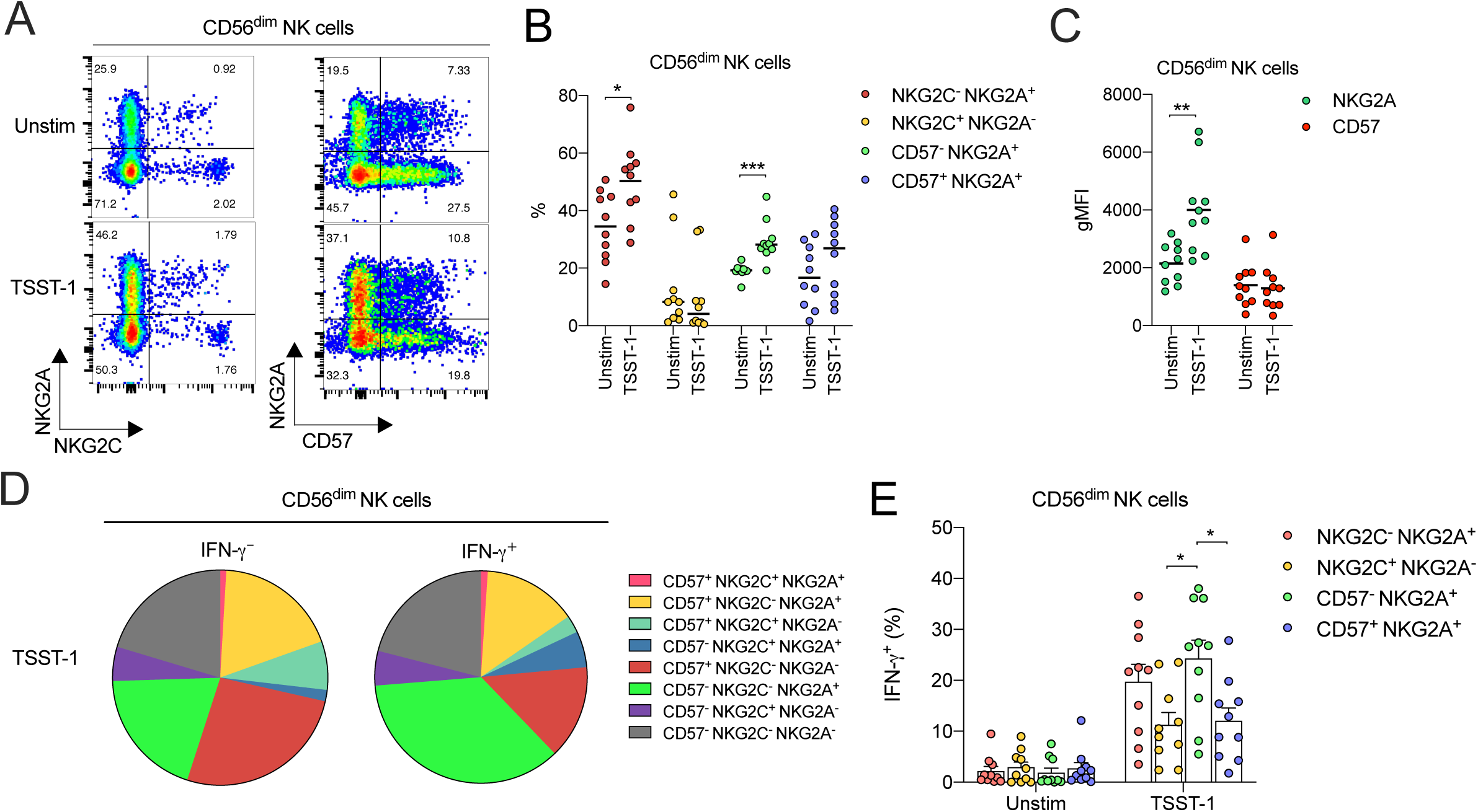
TSST-1 promotes the expansion of CD57^−^ NKG2A^+^ NK cells. (A) Representative flow cytometry plots depicting expression levels of NKG2A vs NKG2C or NKG2A vs CD57 on CD56^dim^ NK cells from human PBMCs collected from healthy controls that were cultured in medium alone (Unstim) or stimulated with TSST-1 for 24h. (B) Scatter dot plot showing the frequency of NKG2C^−^ NKG2A^+^ (red filled circles), NKG2C^+^ NKG2A^−^ (yellow filled circles), CD57^−^ NKG2A^+^ (green filled circles), or CD57^+^ NKG2A^+^ (blue filled circles) subsets among CD56^dim^ NK cells from human PBMCs collected from healthy controls that were cultured in medium alone (Unstim) or stimulated with TSST-1 for 24h. Each dot represents one donor. Line indicates mean value. *p < 0.05, ***p < 0.001; Mann-Whitney U- test. (C) Scatter dot plots depicting geometric mean fluorescence intensity (gMFI) of NKG2A (green filled circles) or CD57 (red filled circles) expression on CD56^dim^ NK cells from human PBMCs collected from healthy controls that were cultured in medium alone (Unstim) or stimulated with TSST-1 for 24h. Each dot represents one donor. Line indicates mean value. **p < 0.01; Mann-Whitney U-test. (D) Pie charts depicting the proportion of IFN-γ^−^ or IFN-γ^+^ CD56^dim^ NK cells that are made up of subsets that express combinations of CD57, NKG2A and/or NKG2C from human PBMCs that were stimulated with TSST-1 for 24h. Data is concatenated from *n =* 6 individual donors. (E) Frequency of IFN-γ^+^ cells among NKG2C^−^ NKG2A^+^ (red filled circles), NKG2C^+^ NKG2A^−^ (yellow filled circles), CD57^−^ NKG2A^+^ (green filled circles), or CD57^+^NKG2A^+^ (blue filled circles) CD56^dim^ NK cells from human PBMCs collected from healthy controls that were cultured in medium alone (Unstim) or stimulated with TSST-1 for 24h. Each dot represents one donor. Data are shown as mean ± SEM; *p < 0.05; One-way ANOVA with Tukey’s post-hoc test.

To explore the effects of SAg stimulation on CD57^−^ NKG2A^+^ CD56^dim^ NK cell subsets further, we examined longer stimulation time periods using TSST-1 as a candidate SAg to determine whether such phenotypic polarization was more pronounced over time. Indeed, the percentage of CD57^−^ NKG2A^+^ or NKG2C^−^ NKG2A^+^ CD56^dim^ NK cells continued to increase significantly following a period of 48h or 72h stimulation (Figures 4A & 4B). In line with these observations, NKG2A expression levels on the surface of CD56^dim^ NK cells were significantly increased further following 48h or 72h stimulation (Figure 4C). Furthermore, CD16 expression levels were significantly downregulated by TSST-1 stimulation on CD56^dim^ NK cells and CD57^−^ NKG2A^+^ subsets, whilst CD56 expression also increased significantly over time with stimulation (Figure 4C & 4D). These changes were consistent with the notion of TSST-1 expanding a less differentiated NK cell phenotype upon stimulation. Additionally, the highest expression frequencies of the proliferation marker Ki67 were seen in CD57^−^ NKG2A^+^ CD56^dim^ NK cells after 72h stimulation with TSST-1 (Figure 4E).

**Figure 4.**
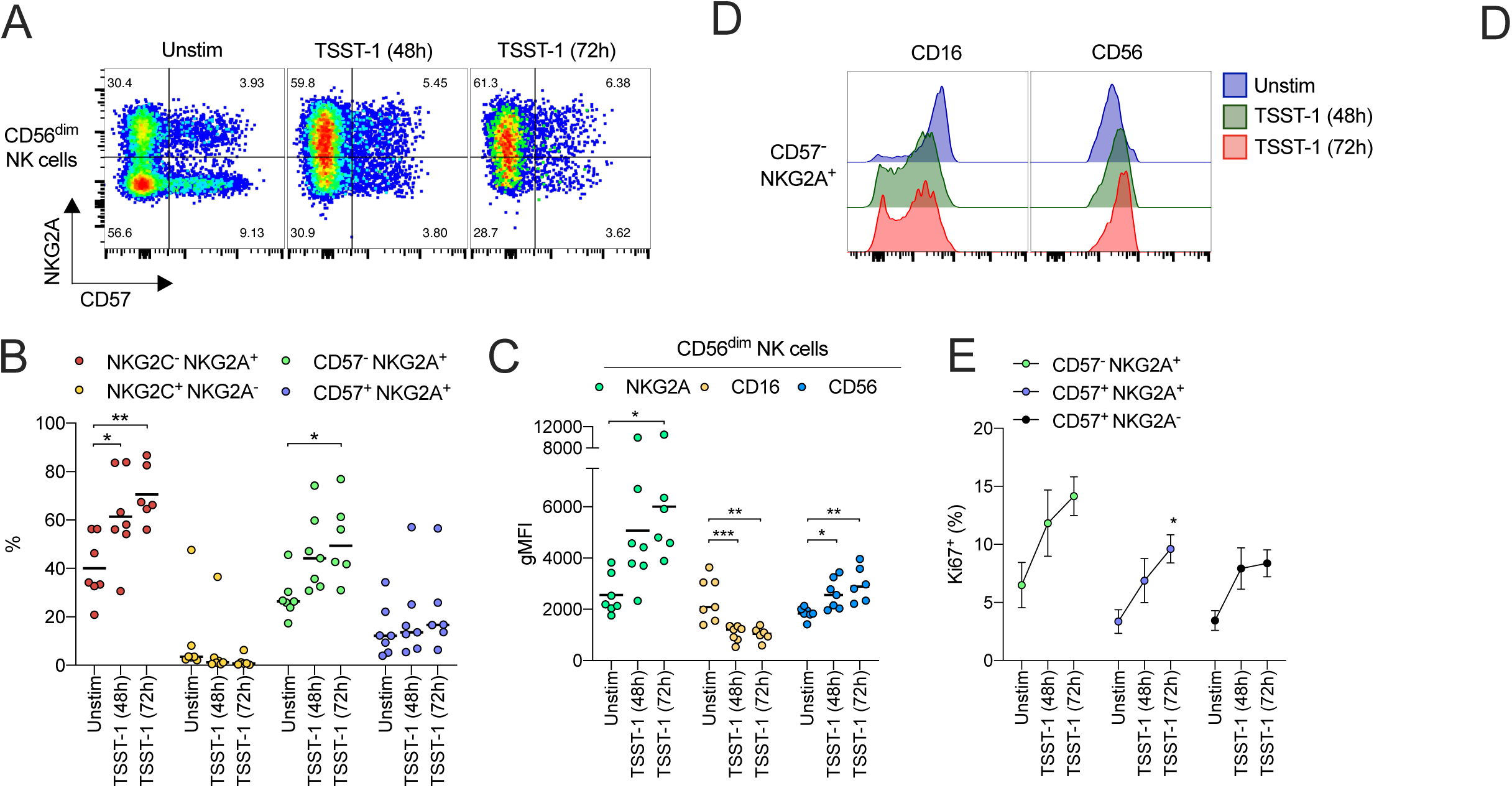
TSST-1-mediated alterations in NKG2A^+^ NK cells are more pronounced over time. (A) Representative flow cytometry plots depicting expression levels of NKG2A vs CD57 on CD56^dim^ NK cells from human PBMCs collected from healthy controls that were cultured in medium alone (Unstim) or stimulated with TSST-1 for 48h or 72h. (B) Scatter dot plots showing the frequency of NKG2C^−^ NKG2A^+^ (red filled circles), NKG2C^+^ NKG2A^−^ (yellow filled circles), CD57^−^ NKG2A^+^ (green filled circles), or CD57^+^ NKG2A^+^ (blue filled circles) subsets among CD56^dim^ NK cells from human PBMCs collected from healthy controls that were cultured in medium alone (Unstim) or stimulated with TSST-1 for 48h or 72h. Each dot represents one donor. Line indicates mean value. *p<0.05, **p < 0.01; One-way ANOVA with Tukey’s post-hoc test. (C) Scatter dot plots depicting geometric mean fluorescence intensity (gMFI) of NKG2A (green filled circles), CD16 (yellow filled circles) or CD56 (blue circles filled circles) expression on CD56^dim^ NK cells from human PBMCs collected from healthy controls that were cultured in medium alone (Unstim) or stimulated with TSST-1 for 48h or 72h. Each dot represents one donor. Line indicates mean value. *p<0.05, **p < 0.01, ***p<0.001; One-way ANOVA with Tukey’s post-hoc test. (D) Representative overlay histograms showing expression of CD16 or CD56 on CD57^−^ NKG2A^+^ CD56^dim^ NK cells from human PBMCs collected from healthy controls that were cultured in medium alone (Unstim) or stimulated with TSST-1 for 48h or 72h. (E) Frequency of Ki67^+^ cells among CD57^−^ NKG2A^+^ (green filled circles), CD57^+^ NKG2A^−^ (blue filled circles) or CD57^+^ NKG2A^−^ (black filled circles), CD56^dim^ NK cells from human PBMCs collected from healthy controls that were cultured in medium (*n =* 7) alone (Unstim) or stimulated with TSST-1 for 48h (*n =* 7) or 72h (*n =* 6). Data are shown as mean ± SEM; *p < 0.05; One-way ANOVA with Tukey’s post-hoc test.

Collectively, these data show that TSST-1 stimulation promotes the expansion of NKG2A^+^ NK cells within human PBMCs, favouring highly proliferative, IFN-γ producing CD57^−^ NKG2A^+^ subsets whilst simultaneously driving a phenotypic shift towards this population.

### TSST-1-mediated promotion of a CD57^-^NKG2A^+^ NK cell phenotype is IL-12- independent

IL-12 is known to upregulate NKG2A expression on NK cells(71, 72). Thus, we next used an anti-IL-12p70 neutralizing antibody to determine if the increased percentage of CD57^−^ NKG2A^+^ CD56^dim^ NK cells elicited after TSST-1 stimulation of PBMCs involved an IL-12- dependent mechanism. As expected, IL-12 blockade reduced the percentage of IFN-γ^+^ CD56^dim^ NK cells elicited after 48h stimulation with TSST-1 (Figures S9A & S9B), agreeing with published data(66). However, neutralization of IL-12 had no discernible effect on the expansion of CD57^−^ NKG2A^+^ CD56^dim^ NK cells that occurred after 48h TSST-1 stimulation (Figures 5A & 5B). TSST-1-mediated changes in the percentages of NKG2C^−^ NKG2A^+^ or CD57^+^ NKG2A^+^ subsets were also similarly unaffected by IL-12 blockade (Figure 5B). In line with these observations, we saw no effect on NKG2A expression changes on the surface of CD56^dim^ NK cells by TSST-1 (Figure 5C). Furthermore, TSST-1-mediated changes in CD16 and CD56 expression were also IL-12-independent (Figure 5C). These data suggest that whilst IL-12 is an important factor in SAg-induced IFN-γ production from CD56^dim^ NK cells, it is not involved in driving CD56^dim^ NK cells towards a CD57^−^ NKG2A^+^ NK cell phenotype.

**Figure 5.**
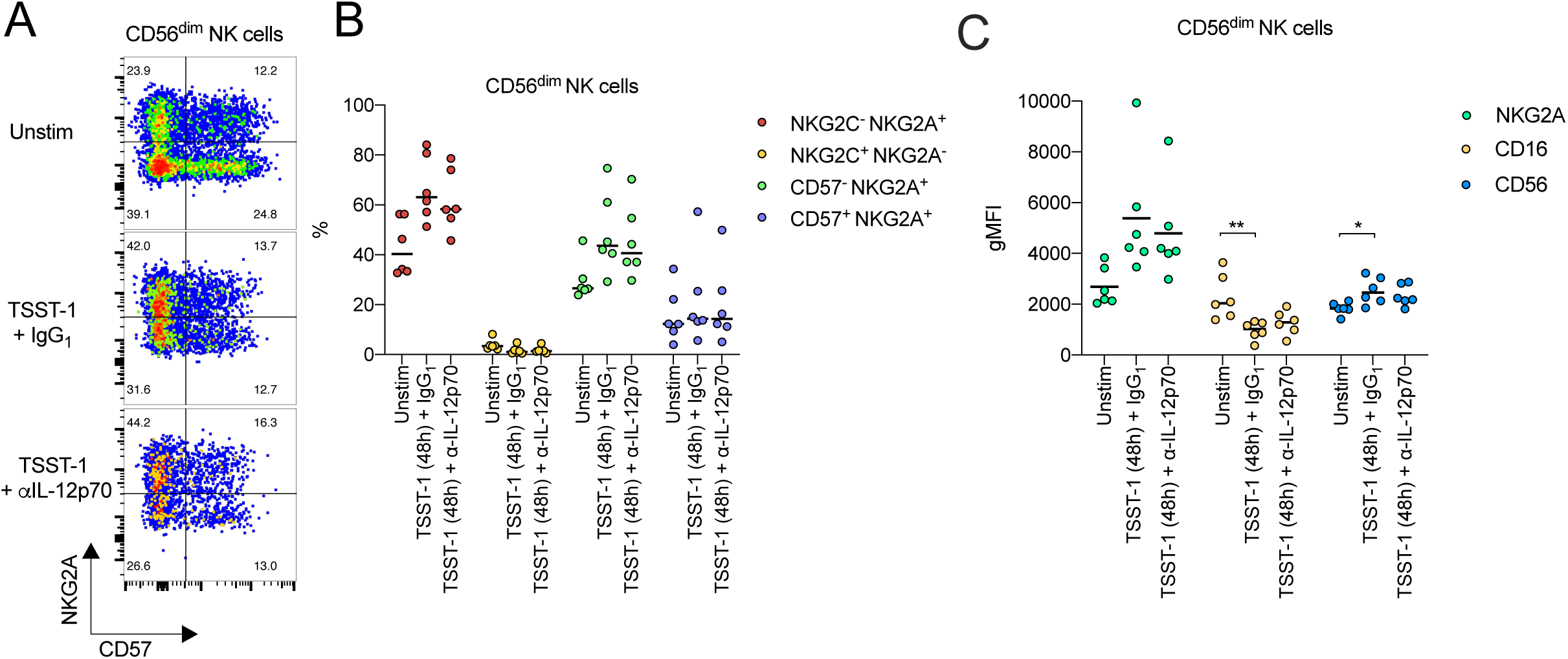
TSST-1-mediated polarisation towards a CD57^−^ NKG2A^+^ NK cell phenotype involves an IL-12-independent mechanism. (A) Representative flow cytometry plots depicting expression levels of NKG2A vs CD57 on CD56^dim^ NK cells from human PBMCs collected from healthy controls that were cultured in medium alone (Unstim) or pretreated with anti-human IL-12p70 or an isotype control antibody (IgG_1_) before being stimulated with TSST-1 for 48h. (B) Scatter dot plot showing the frequency of NKG2C^−^ NKG2A^+^ (red filled circles), NKG2C^+^ NKG2A^-^ (yellow filled circles), CD57^−^ NKG2A^+^ (green filled circles), or CD57^+^ NKG2A^+^ (blue filled circles) subsets among CD56^dim^ NK cells from human PBMCs collected from healthy controls that were cultured in medium alone (Unstim) or pretreated with anti-human IL-12p70 or an isotype control antibody (IgG_1_) before being stimulated with TSST-1 for 48h. Each dot represents one donor. Line indicates mean value. (C) Scatter dot plots depicting geometric mean fluorescence intensity (gMFI) of NKG2A (green filled circles), CD16 (yellow filled circles) or CD56 (blue circles filled circles) expression on CD56^dim^ NK cells from human PBMCs collected from healthy controls that were cultured in medium alone (Unstim) or pretreated with anti-human IL-12p70 or an isotype control antibody (IgG_1_) before being stimulated with TSST-1 for 48h. Each dot represents one donor. Line indicates mean value. *p < 0.05, **p < 0.01; One-way ANOVA with Tukey’s post-hoc test.

### CD56^dim^ NK cells in human patients with *S. aureus* bacteraemia are more polyfunctional

Peripheral blood NKG2A^+^ NK cells are effective cytokine producers and differentiation towards CD57^+^ NKG2C^+^ populations generates NK cells that are less cytokine responsive but with more specialized functions such as receptor-driven cytotoxicity(32, 36–38). Furthermore, the co-inhibitory receptor TIGIT can engage target ligands such as CD155(73–76) and the TIGIT-CD155 axis can drive NK cells towards enhanced effector degranulation and cytokine production, notably against CD155^-^ target cells (82). Our data demonstrates that NKG2A^+^ NK cells expand during *S. aureus* bacteraemia and that a positive correlation exists between NK cell activation and TIGIT positivity on NK cells in these patients. As such, we hypothesized that *S. aureus* bacteraemia may generate NK cells with more degranulating and cytokine-producing capacity. To test this, we stimulated PBMCs collected from healthy controls and patients with *S. aureus* or *E. coli* bacteraemia (at hospital admission) with PMA/ionomycin (CD155-independent stimuli) and measured the percentage of CD56^dim^ NK cells that degranulated (i.e. based on surface mobilization of CD107a(^83^)) and produced IFN-γ and/or TNF-α. We found that PMA/ionomycin stimulation yielded a significant increase in CD107a^+^ CD56^dim^ NK cells from patients with *S. aureus* bacteraemia, compared to healthy controls (Figure 6A). In contrast, patients with *E. coli* bacteraemia had lower levels of CD107a^+^ cells compared to that seen in *S. aureus* bacteraemia patients. A high proportion of CD107a^+^ NK cells in *S. aureus* patients were IFN-γ^+^ (Figure 6B) and these CD107a^+^ IFN-γ^+^ NK cells were at significantly higher levels than those found in healthy controls (Figure 6C). This suggested that patients with *S. aureus* bacteraemia may possess more polyfunctional NK cells, compared to healthy controls or patients with *E. coli* bacteraemia. Indeed, using Boolean gating and Simplified Presentation of Incredibly Complex Evaluations (SPICE) software(84) to assess the degree of NK cell polyfunctionality, patients with *S. aureus* bacteraemia displayed a higher proportion of polyfunctional CD56^dim^ NK cells, compared to healthy controls or patients with *E. coli* bacteraemia (Figure 6D). Furthermore, these *S. aureus* patients had fewer unresponsive CD56^dim^ NK cells (CD107a^−^ IFN-γ^−^ TNF-α^−^) compared to healthy controls or patients with *E. coli* bacteraemia (Figure 6D).

**Figure 6.**
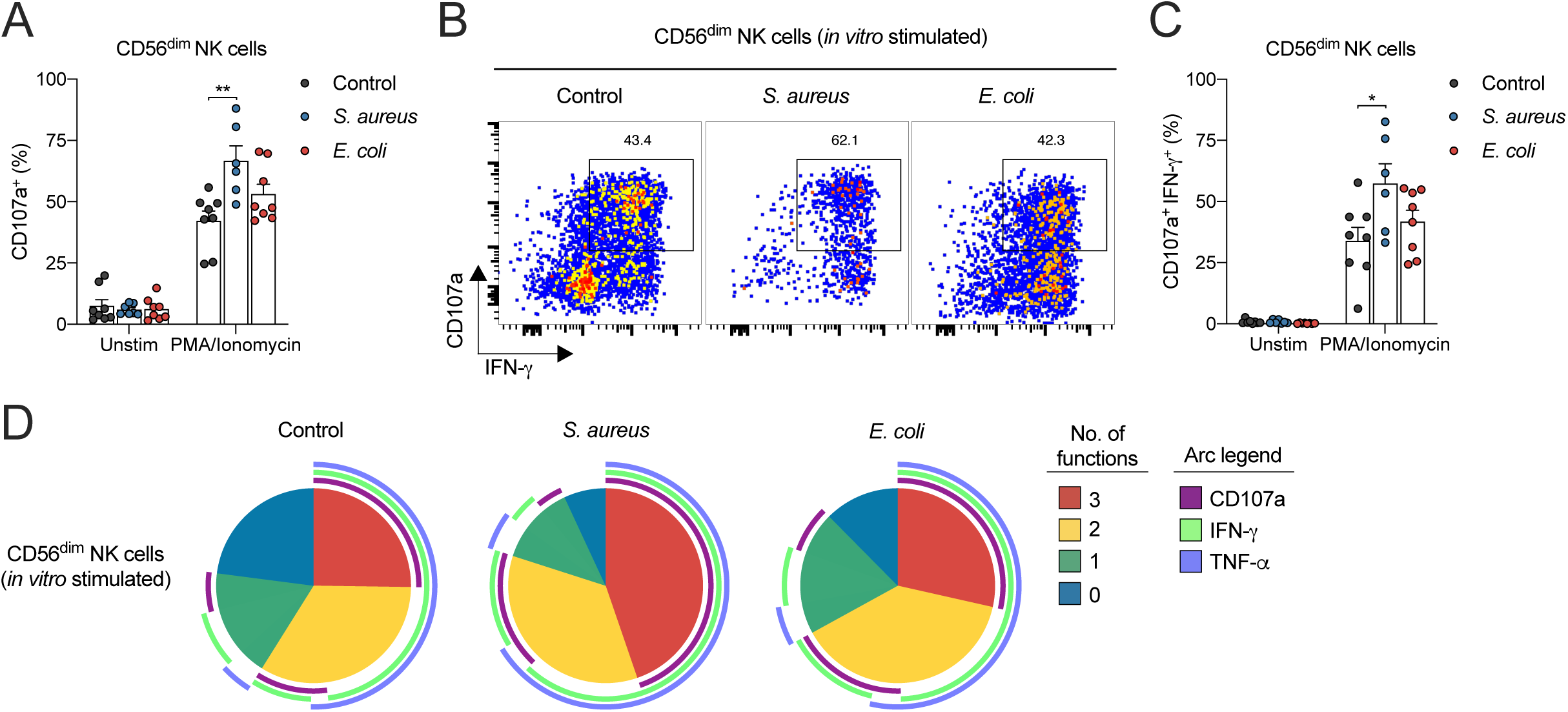
CD56^dim^ NK cells in human patients with *S. aureus* bacteraemia are more polyfunctional, yielding more effector cytokines upon stimulation. (A) Frequency of CD107a^+^ cells among CD56^dim^ NK cells from human PBMCs collected from healthy controls (black filled circles) and patients hospitalized with *S. aureus* (blue filled circles) or *E. coli* (red filled circles) bacteraemia cultured in medium alone (Unstim) or with PMA/Ionomycin for 6h. Data are shown as mean ± SEM; **p < 0.01, One-way ANOVA with Tukey’s post-hoc test. (B) Representative flow cytometry plots depicting expression levels of CD107a and IFN-γ among CD56^dim^ NK cells from human PBMCs collected from healthy controls and patients hospitalized with *S. aureus* or *E. coli* bacteraemia cultured in medium alone (Unstim) or with PMA/Ionomycin for 6h. (C) Frequency of CD107a^+^ IFN-γ^+^ cells among CD56^dim^ NK cells from human PBMCs collected from healthy controls (black filled circles) and patients hospitalized with *S. aureus* (blue filled circles) or *E. coli* (red filled circles) bacteraemia cultured in medium alone (Unstim) or with PMA/Ionomycin for 6h. Data are shown as mean ± SEM; *p < 0.05, One-way ANOVA with Tukey’s post-hoc test. (D) Functional profiles of CD56^dim^ NK cells from human PBMCs stimulated with PMA/Ionomycin for 6h. Pie chart segments from concatenated data (n = 7 - 8) represent the fraction of cells displaying the indicated number of functions (key). Arcs denote individual functions (key).

Collectively, these results demonstrate that CD56^dim^ NK cells in patients with *S. aureus* bacteraemia are functionally more responsive to stimulation, yielding more polyfunctional NK cells, compared to healthy controls and those hospitalized with *E. coli* bacteraemia.

## DISCUSSION

In this study, we observed an increase in the frequency of NKG2A^+^ NK cells in patients with *S. aureus* bacteraemia compared to healthy controls and found that NK cells from these patients were functionally more responsive to stimulation, displaying evidence of being more polyfunctional and, thus, being capable of producing more effector cytokines, such as IFN-γ. Accordingly, our data suggest that systemic *S. aureus* infection in humans promotes a phenotypic shift towards NK cell subsets with greater IFN-γ producing capacity and notionally agrees with the concept that NK cells may promote inflammation-mediated disease pathogenesis during *S. aureus* bacteraemia. As far as we are aware, our study is the first to demonstrate this phenotypic alteration towards NKG2A^+^ NK cells in this patient group and is consistent with studies in mouse models of *S. aureus* infection suggesting that NK cell-released IFN-γ may promote immune pathogenesis. In contrast, we did not observe the same alteration of NKG2A^+^ NK cells in patients with *E. coli* bacteraemia, despite mouse models indicating that NK cells may also be immunopathogenic during systemic *E. coli* infection(57). This suggests that *S. aureus* itself may be directly responsible for triggering this change in the frequency of NKG2A^+^ NK cells in patients. Indeed, our data shows that SAgs from *S. aureus*, namely TSST-1, can upregulate NKG2A expression on NK cells and promote the expansion of highly proliferative, IFN-γ producing CD57^−^ NKG2A^+^ subsets.

SAgs may not be the sole driver behind these phenotypic changes. *S. aureus* also encodes pore-forming toxins that affect NK cell behaviour, however, their effect on NKG2A expression is unknown and they are reported to deplete their frequencies(67) or inhibit their IFN-γ producing capacity(68). Beyond this, it is possible that unique features of the immune response to *S. aureus* may promote this increased frequency of NKG2A^+^ NK cells. IL-12 can upregulate NKG2A expression on NK cells(71, 72) yet we found that plasma IL-12p70 levels were lower in patients with *S. aureus* bacteraemia, compared to healthy controls or patients with *E. coli* bacteraemia. Transforming growth factor-β (TGF-β) is capable of increasing NKG2A expression on T cells(85) and TGF-β activation occurs in mouse models of *S. aureus* infection, principally through the actions of α-hemolysin (86). Thus, the increased frequency of NKG2A^+^ NK cells could be mediated by TGF-β, although our observed reduction of NKG2C^−^ NKG2A^+^ CD56^+^ T cells in patients with *S. aureus* bacteraemia compared to healthy controls, would not be consistent with this. Nevertheless, expanding our understanding of how *S. aureus* modulates NK cell activity is important to determine whether these effects are exclusive to *S. aureus* or are observed during infection with other bacterial pathogens.

In conjunction with observing an increase in the frequency of NKG2A^+^ NK cells in patients with *S. aureus* bacteraemia, we also detected a concurrent reduction in the percentage of CD57^+^ NKG2C^+^ NKp30^−^ Siglec-7^−^ NK cells compared to healthy controls and patients with *E. coli* bacteraemia. This phenotype has been used to define highly differentiated, adaptive-like NK cells that are proposed to contribute to antiviral protection and likely represent a host response to HCMV-encoded immune evasion (46, 47, 77, 78). For instance, adaptive-like NK cells are also ILT2/LIR1^+^ and CD2^high^ (77, 78, 87), both of which are pathways targeted by the HCMV genes, UL18 and UL148 respectively(88, 89). The NKG2A/NKG2C axis represents another. NKG2C, in complex with C-type lectin CD94, binds to, and is activated by, HLA-E(90), a non-classical major histocompatibility complex class I molecule that can present self and viral peptides, including polymorphic peptides derived from the HCMV-encoded UL40 protein to NK cells(91). However, NKG2A, which similarly dimerizes with CD94, also binds HLA-E but with greater affinity(90, 92), but delivers an inhibitory signal to NKG2A^+^ NK cells (93, 94). In line with these concepts, it is plausible that *S. aureus* may also modulate HLA-E expression levels to evoke changes in the frequency of NKG2C^−^ NKG2A^+^ NK cell subsets. SAgs may also reduce the expression of HLA-E on different cell types including APCs releasing its inhibitory effect on NKG2A(95), and leading to greater activation of NKG2A^+^ NK cell subsets and subsequent expansion of CD57^−^ NKG2A^+^ NK cell populations. The reduction in the percentage of CD57^+^ NKG2C^+^ NKp30^-^ Siglec-7^-^ NK cells may also be reflective of these cells being converted to NKG2A^+^ NK cells, which also increased in frequency within NKp30^-^ Siglec-7^-^ NK cell populations. Further work is required to determine if this is a true feature of *S. aureus* infection but also how, and if, severe bacterial infections may affect adaptive-like NK cell populations.

The ability for SAgs from *S. aureus*, including TSST-1, to alter NK cell functionality has been reported before, notably in how they indirectly induce NK cells to produce high levels of IFN-γ through an IL-12-dependent mechanism (65, 66). Whilst our data agree with these observations, we discovered that SAgs, notably TSST-1, promoted the expansion of highly proliferative, IFN-γ producing CD57^−^ NKG2A^+^ NK cells with reduced CD16 expression, but, this process was not dependent on IL-12 even though SAgs promote IL-12 production from myeloid cells (66, 79, 80) and IL-12 itself can upregulate NKG2A expression on NK cells(71, 72). Consequently, the core mechanism involved requires elucidation. TSST-1 has been shown to increase the percentage of NKG2A^+^ T cells in time-course stimulation assays where PBMCs were depleted of CD94^+^ cells (96). Furthermore, whilst these effects were relatively modest, even after 8-10 days of stimulation, co-culture with IL-15 or TGF-β amplified this process significantly(96, 97). This suggested that TSST-1 was effective at converting NKG2A^−^ T cells into NKG2A^+^ T cells in tandem with IL-15 or TGF-β. Furthermore, IL-12 supplementation had no effect on this process(96). As such, our data on SAg-mediated changes in NK cells may reflect NKG2C^−^ NKG2A^−^ NK cells converting into NKG2C^−^ NKG2A^+^ NK cell subsets, rather than specific proliferation events occurring solely in pre-existing NKG2C^−^ NKG2A^+^ NK cells. However, it is worth noting that these T cell-based studies did not explore what factors were critical for TSST-1 being able to increase the percentage of NKG2A^+^ T cells in the absence of supplemented co-factors such as IL-15 or TGF-β. Recently, it was shown that SEB-induced changes in NKG2A expression on CD8^+^ T cells were impaired by CD40 blockade, which also prevented IL-12 production(98). Whilst it is still unknown if CD40 influences SAg-mediated changes in NKG2A expression on NK cells or the expansion of CD57^−^ NKG2A^+^ NK cells seen in our experiments, it has been reported that SAgs can bind to CD40(26), which is found on APCs, and such binding could promote these phenotypic changes. Nonetheless, it will be important to dissect the critical factors that enable SAgs to increase the frequency of CD57^−^ NKG2A^+^ NK cells and compare such dependency patterns in T cells, where direct binding by SAgs to TCRs occurs. To this end, the notion that SAgs may directly bind NK cells(99) has been refuted since human NK cells alone appear to be unresponsive to SAgs, only occurring when T cells and monocytes are co-cultured with them (66).

Compared to what is known about NK cell-mediated immunity to viruses, our knowledge about how NK cells respond to bacterial infections is very limited and is complicated by the ambiguous nature of their precise role. Indeed, it appears that there are species-specific differences in the control and immunopathology of bacterial infections by NK cells. Our data on the NK cell response to *S. aureus* bacteraemia supports the concept that NK cells can promote immune pathology during bacterial infection and that this is linked to the impact of NK cell-derived production of IFN-γ, and likely other cytokines, in driving excessive inflammation. However, more evidence is needed to examine the precise role of NK cells across different types of bacterial infection in humans. Indeed, our study had some limitations, notably in the modest number of patients with *S. aureus* or *E. coli* bacteraemia that we enrolled but also that they were predominantly (for *E. coli*) or all (for *S. aureus*) female. Thus, larger longitudinal studies with better sex matching are needed to expand this further and learn more about NK cell-mediated inflammation to systemic *S. aureus* infection. This is perhaps even more relevant given the increased mortality risk seen in female patients with *S. aureus* bacteraemia(100, 101), which ultimately means a better understanding of sex-specific nuances in NK cell-mediated immunity is even more crucial. Furthermore, it will be important to compare findings with data collected from patients with bloodstream infections caused by other bacterial organisms.

In summary, we observed an increase in polyfunctional CD56^dim^ NK cells with greater cytokine producing potential in patients with *S. aureus* bacteraemia. These patients also exhibited increased frequencies of NKG2A^+^ NK cell subsets, whilst SAgs from *S. aureus* promoted the expansion of IFN-γ producing, CD57^−^ NKG2A^+^ NK cells through an IL-12- independent mechanism. As such, our data supports the concept that immune pathogenesis during *S. aureus* infection may be linked to over-production of pro-inflammatory cytokines and that alterations in NK cell-released IFN-γ could be a driver of immune pathogenesis during *S. aureus* infection. These mechanisms require further elucidation but imply that therapeutic approaches combining immune modulatory agents with antibiotics need to be explored.

## MATERIALS AND METHODS

### Subjects and ethical approval

The recruitment of adult patients admitted to the University Hospital of Wales with invasive bloodstream *S. aureus* or *E. coli* infections (bacteraemia) was approved by the NHS Research Ethics Committee (reference 21/WA/0202, IRAS project ID 277598) and was sponsored by Cardiff University and Cardiff & Vale University Health Board. All participants provided written informed consent for the collection of peripheral blood samples and subsequent analyses in accordance with the principles of the Declaration of Helsinki. Patients were excluded if they were pregnant, breastfeeding or females of childbearing age in whom a pregnancy test had not been performed. Furthermore, participants considered too unwell, with limited life expectancy, with mental incapacity or language barriers which preclude adequate understanding or those with an estimated glomerular filtration rate <45 ml/min/1.73m^2^ where excluded. The study cohort comprised patients with microbiologically confirmed *S. aureus* (age ranging from 27 to 79 years (median 63 years)) or *E. coli* (age ranging from 31 to 74 years (median 68 years)) bacteraemia. Peripheral venous blood samples were collected from bacteraemia patients at hospital admission (within 2-6 days post-diagnosis) from patients with *S. aureus* (*n* = 7; median 3 days post-diagnosis) or *E. coli* (*n* = 8; median 3 days post-diagnosis) bacteraemia. Matched donor blood samples collected at convalescence (as close to 30 days post-hospital discharge as possible) were collected from *n* = 4 patients with *S. aureus* bacteraemia and *n* = 8 patients with *E. coli* bacteraemia. Healthy control blood donors (*n* = 8; age range 59–72 years, median 66 years) were recruited as controls for the study if there was no history of recent (<1 year) bloodstream *S. aureus* or *E. coli* infections, were over the age of 18, had the capacity to consent or in the event of temporarily reduced capacity, consent from the next-of-kin (or another suitable consultee) was gained. Peripheral venous blood samples were collected from healthy controls (*n* = 8) at point of enrolment. For experimental assays not involving bacteraemia patients and study controls (i.e. experiments involving *in vitro* SAg stimulation assays), peripheral venous blood samples were collected from healthy volunteers and buffy coats purchased from the Welsh Blood Service. The use of venous blood samples from healthy volunteers was approved by the Cardiff University School of Medicine Research Ethics Committee (18/4). Written informed consent was obtained from all donors in accordance with the principles of the Declaration of Helsinki.

### Cell culture and *in vitro* stimulation assays

Human peripheral blood mononuclear cells (PBMCs) were isolated via density gradient centrifugation using Histopaque-1077 (Sigma-Aldrich) and cryopreserved in foetal bovine serum containing 10% dimethyl sulfoxide (Sigma-Aldrich). Human PBMCs were thawed and rested in RPMI 1640 medium supplemented with 10% fetal bovine serum, 100 U/mL penicillin, 100 μg/mL streptomycin, and 2 mM L-glutamine (all from Thermo Fisher Scientific) (R10) at 37°C and in 5% CO_2_ prior to being seeded in 24-well or 96-well culture plates for *in vitro* stimulation assays. Cell Activation Cocktail (BioLegend) was used to deliver 25 ng/ml phorbol-12-myristate 13-acetate (PMA) and 500 ng/ml ionomycin to cultured PBMCs for 6h at 37°C. Recombinant forms of SEA, SEB or TSST-1 from *S. aureus* (Toxin Technology), each at a dose of 100 ng/mL to align with previous work(19, 102–104), were used to stimulate PBMCs for 24h, unless otherwise stated. Unstimulated cells were used as negative controls. In mechanistic experiments, human PBMCs were preincubated for 30 min with 2 μg/mL anti-human IL-12 (clone 24910; BioTechne) or 2 μg/mL mouse IgG_1_ (clone MOPC-21; BioLegend). In functional experiments, anti-CD107a–BV421 (clone H4A3; BioLegend) was added to the cultures with PMA/ionomycin or each individual SAg(105), and protein transport was blocked for the final 6h with GolgiPlug (1:1,000; BD Biosciences) and GolgiStop (1:1,500; BD Biosciences).

### Flow cytometry

Cells were washed in Dulbecco’s phosphate-buffered saline (Thermo Fisher Scientific), labelled for 15–30 min at room temperature with Zombie Aqua (BioLegend) for live/dead cell exclusion. Surface stains were performed for 30 min at 4°C using combinations of the following directly conjugated monoclonal antibodies: (i) anti-CD3–APC/Fire 750 (clone SK7), anti-CD3–PerCP (clone SK7), anti-CD14–APC/Fire 750 (clone M5E2), anti-CD16– BV711 (clone 3G8), anti-CD56–BV605 (clone HCD56), anti-CD56–BV711 (clone HCD56), anti-CD57–PE/Dazzle 594 (clone QA17A04), anti-CD69–APC (clone FN50), anti-CD69– BV421 (clone FN50, anti-NKG2C–PE (clone S19005E), anti-NKp30–APC/Fire 750 (clone P30-15), anti-Siglec-7–APC/Fire 750 (clone 6-434) and anti-TIGIT-BV421 (clone A15153G) from BioLegend; (ii) anti-CD14 V500 (clone M5E2) and anti-CD19 V500 (clone HIB19) from BD Biosciences; and (iii) anti-NKG2A-VioBright FITC (clone REA110) from Miltenyi Biotec. Cytosolic/intranuclear expression of Ki67 was detected using anti-Ki67–BV421 (clone B56; BD Biosciences) in conjunction with a FOXP3 Transcription Factor Staining Buffer Kit (Thermo Fisher Scientific). Intracellular cytokines were exposed using a Cytofix/Cytoperm Plus Fixation/Permeabilization Solution Kit (BD Biosciences) or FOXP3 Transcription Factor Staining Buffer Kit (Thermo Fisher Scientific) and stained for 30 min at 4 °C with combinations of the following directly conjugated monoclonal antibodies: anti-IFN-γ–APC (clone B27), anti-IFN-γ–PE/Cyanine7 (clone B27) and anti-TNF-α–APC (clone MAb11) from BioLegend. All flow cytometry panels were validated using individually stained anti-mouse Ig, 1/Negative Control Particles (BD Biosciences) or MACS anti-REA Comp Bead kit (Miltenyi Biotec). Data were acquired using an Attune NxT flow cytometer (Thermo Fisher Scientific) and analyzed using FlowJo software version 10.10 (FlowJo LLC).

### IL-12p70-specific ELISA

Plasma was collected from patients with *S. aureus* or *E. coli* bacteraemia and healthy controls during the density gradient centrifugation of peripheral blood using Histopaque-1077. Aliquots were prepared and stored at −80 °C until further use. IL-12p70 levels were quantified using a commercial ELISA kit (Mabtech). Nunc MaxiSorp 96-well plates (BioLegend) were coated with an anti-IL-12p40 capture antibody (2 μg/ml; Mabtech clone MT86/221) and incubated overnight at 4°C. Following incubation, plates were blocked with PBS supplemented with 0.1% bovine serum albumin and 0.05% Tween-20 (Sigma Aldrich)) for 1h at room temperature (RT) to reduce non-specific binding. Following this, coated plates were washed three times with PBS containing 0.05% Tween-20 before undiluted plasma samples and recombinant IL-12p70 protein (serially diluted from 1 ng/ml after preparation from a stock concentration of 1 μg/ml to prepare a standard curve) were added in duplicate and incubated for 2h at RT. After further washing, plates were incubated for 1h at RT with a biotin conjugated anti-IL-12p70 detection antibody (1 μg/ml; Mabtech clone MT704) diluted to 1 μg/ml before the process was repeated using streptavidin conjugated to alkaline phosphatase (Mabtech; 1:1000 dilution of stock protein). Following further washing with PBS containing 0.05% Tween-20, Para-nitrophenylphosphate (Mabtech) was added to each well and incubated for 60 mins at RT to develop in the absence of light. Optical densities were read at 405nm using the CLARIOstar microplate reader (BMG Labtech). Samples were blinded to eliminate operator bias. Standard curves were plotted using a nonlinear regression model on Prism (GraphPad v10.1.0), with concentrations calculated in reference to the curve. All recombinant IL-12p70 protein standard and human plasma sample aliquots were thawed immediately prior to use.

### Statistics

Differences between two groups were evaluated using a Mann-Whitney U-test whilst differences between three or more groups were evaluated using a one-way ANOVA with Tukey’s post-hoc test in Prism version 8.3.1 (GraphPad). Analysis of paired samples was performed using Wilcoxon matched-pairs sign ranked test whilst tests of correlation between two variables were performed using Spearman’s rank correlation test. Significance was assigned at p<0.05 and depicted in figures with asterisk where achieved.

## ACKNOWLEDGEMENTS

Figure 1A was created using BioRender.com.

## AUTHOR CONTRIBUTIONS

KD: Investigation, Formal Analysis, Resources, Validation, Visualization Writing – Review & Editing

AR: Investigation, Formal Analysis

SK: Conceptualization, Methodology, Writing – Review & Editing

ECYW: Conceptualization, Methodology, Writing – Review & Editing

ME: Supervision, Resources, Writing – Review & Editing

JU: Conceptualization, Funding Acquisition, Project Administration, Resources, Writing – Review & Editing

JEM: Conceptualization, Formal Analysis, Funding Acquisition, Investigation, Methodology, Project Administration, Resources, Supervision, Visualization, Writing – Original Draft Preparation, Writing – Review & Editing

